# Species-level identification of the critically endangered European sturgeon *Acipencer sturio* using DNA-based approaches

**DOI:** 10.1101/2023.05.05.539572

**Authors:** Emilie Chancerel, Marie-Laure Acolas, Olivier Lepais

**Affiliations:** Univ. Bordeaux, INRAE, BIOGECO, Plateforme Génome Transcriptome de Bordeaux, 33610 Cestas, France; INRAE, UR EABX, 50 Avenue de Verdun, 33612 Gazinet-Cestas, France

## Abstract

The native European sturgeon *Acipenser sturio* is the focus of restoration programs. However, *A. sturio* identification by direct observation is not always obvious for non-specialists. We aimed to develop a quick and easy genetic tool applicable to identify the species at risk in case of incidental capture or monitoring either at sea or in the watershed. We showed that primers and probe developed allow specific amplification for *A. sturio* by PCR, qPCR and Real-Time Loop-mediated isothermal amplification (LAMP) techniques. We determine benefits and limits of each method and highlight perspectives for field applications within the frame of conservation programs.

## Introduction

Sturgeon species have a worrisome status all other the world (Billard and Lecointre 2000). In Western Europe the native European sturgeon *Acipenser sturio* is critically endangered (Gessner *et al*. 2022) and restoration programs were implemented in France (Adam *et al*. 2020) and Germany (Gessner *et al*. 2010) with regular stocking in the Gironde Garonne Dordogne (Roques *et al*. 2018) and in the Elbe. However, many non-native sturgeon species can be found in the watersheds due to incidental release from hatchery farms. Since the *A. sturio* species is fully protected and the identification by direct observation is not always obvious for non-specialists, a quick and easy distinction with a reliable genetic tool in the field could be a valuable development to identify the species at risk in case of incidental capture or monitoring either at sea or in the watershed.

The purpose of this study was to discriminate easily *A. sturio* from other more or less closely related *Acipenser* species (Krieger *et al*. 2008) using DNA-based approaches. We focused on species present in hatcheries in Western Europe such as: *A. gueldenstaedtii, A. stellatus, A. transmontanus, A. baerii, A. persicus, Huso huso*, and we also add the genetically and morphologically closest species to *A. sturio* which is *A. oxyrinchus* (Krieger *et al*. 2008). We compared different amplification techniques including PCR, qPCR and Real-Time Loop-mediated isothermal amplification (LAMP) to identify the most easily applicable for routine testing.

## Materials and methods

### Fish samples and DNA isolation

Fin of *A. sturio, A. gueldenstaedtii, A. stellatus, A. transmontanus, A. baerii, A. persicus, A. oxyrinchus* and *Huso huso* reared in captivity have been collected and preserved in 90% ethanol. Pieces of fin (2 mm x 2 mm) were used to extract DNA using DNeasy Blood & Tissue kit (Qiagen, Hilden, Germany). DNA was quantified using Qubit fluorimeter with Qubit BR Assay Kit.

### Design of *A. sturio* specific primers for PCR and qPCR assays

Specific *A. sturio* primers for PCR and qPCR were designed using IDT Primer Quest Tool and aligned using Clustal W (Larkin *et al*. 2007) to cytochrome b gene sequences from *A. sturio, A. gueldenstaedtii, A. stellatus, A. transmontanus, A. baerii, A. persicus, H. huso* obtained from GenBank (Supporting Material Table S1). The designed primers Cytb_AS9_F: AGCCCTAGTGGCCGATATATTA and Cytb_AS9_R: TTTCCAGTCAGCCCGTTATAG target sequences that are specific to *A. sturio* and differ by several substitutions from the other species (Supporting Material Figure S1). The primers are expected to amplify a 142 bp fragment. A specific PrimeTime qPCR probe (IDT) was also designed (Cytb_AS9_probe: FAM-TTGGACAAGTGGCTTCCACAGTCT).

### Design of *A. sturio* specific primers for LAMP assays

An alignment of cytochrome c oxidase subunit I (COI) gene sequences from *Acipenser sturio, A. gueldenstaedtii, A. stellatus, A. transmontanus, A. baerii, A. persicus* and *Huso huso* (Supporting Material Table S2) was performed using Clustal W (Larkin *et al*. 2007) program to find polymorphic regions between species. Clustal Omega (Sievers *et al*. 2011) and Morphocatcher (Shirshikov *et al*. 2019) generate a fasta file by species specifying the detected mutations and PrimerExplorer v5 program (Eiken Chemical Co. LTD, Tokyo, Japan; http://primerexplorer.jp/e) was used on the *A. sturio* fasta file for primer designs (Option “Specific design”; primer length between 18 and 22 bp). The design with maximum mismatch on primers F3/B3 and FIP/BIP and lowest deltaG dimer was retained (Supporting material Table S2 and Figure S1); F3: CTTTTGGCCTCCTCTGGC, B3: GTCTGATACTGGGATACTGCG, FIP: AGGCTCCTGCATGGGCTAGGTTTTTTCTGGCACAGGATGAACCG and BIP: CCTTCACCTGGCTGGGGTATCTTTTTTGGGGGCTTCATGTTAATGATTG.

### PCR amplification

The set of specific *A. sturio* primers Cytb_AS9_F/Cytb_AS9_R was used combined with another primer pair specific to fish (Xiong *et al*. 2022) as a positive control for PCR success. Two pairs of fish primers were tested independently with specific *A. sturio* primers: L14735/H15149R (AAAAACCACCGTTGTTATTCAACTA / GCDCCTCARAATGAYATTTGTCCTCA ; 403 bp) and L14841/H15149 (AAAAAGCTTCCATCCAACATCTCAGCATGATGAAA / AAACTGCAGCCCCTCAGAATGATATTTGTCCTCA ; 348 bp). Amplifications were performed in a 20 µL reaction volume with either 2 ng or 20 ng of input DNA from the eight studied species, 4 µL of 5X HotFirepol Blend Master Mix (Solis Biodyne, Estonia), 625 nM of L14735/H15149R or L14841/H15149 and Cytb_AS9_F/Cytb_AS9_R. One negative control (water instead of DNA) was included in the PCR run to control for any cross-contamination.

The PCR cycling protocol consisted of the following steps: 95°C for 12 min; 30 cycles of 95 °C for 45 s ; 56°C for 45 s, 72 °C for 45 s and a final elongation at 72 °C for 5 min. The reaction products were visualized under UV light after electrophoresis of 3 μL of the reaction solution combined with 3 µL of 6x DNA Loading Dye Buffer (Solis Biodyne) in a 3% agarose gel electrophoresis (100 Volts / 30 min). The fish marker is used to verify that the amplification is successful for each sample: for all fish species except *A. sturio* an unique band of the expected size fish marker (403 bp or 348 bp) should appear on the gel ; for *A. sturio* two bands should appear on the gel, one at the expected size fish marker (403 bp or 308 bp) and another one at the expected size species marker (142 bp); the negative control should not show any band.

### qPCR amplification

Real-time amplifications were performed using 1 ng of input DNA from the eight studied species, 4 µL of 5X SolisFAST® Probe qPCR (no ROX; Solis Biodyne), 400 nM of Cytb_AS9_F/ Cytb_AS9_R primers and 150 nM of CytB_AS9 probe in a total volume of 20 µL. qPCR reactions were performed in triplicates for each sample on a LightCycler 480 (Roche) and one qPCR reaction per sample was tested on Franklin™ Real-Time PCR Thermocycler (Biomeme). One negative control (water instead of DNA) was included in qPCR runs to control for cross-contamination. Successive DNA dilutions from 10 ng down to 1 fg were used as DNA template to determine the sensitivity of detection. qPCR runs were performed with the following cycle parameter: 95°C for 3 min, 40 cycles of 95°C for 5 s and 63°C for 20 s. qPCR products were evaluated based on C_t_ values.

### Real time LAMP amplification

LAMP was performed in a 20 µL volume containing 12 µL of GspSSD2.0 Isothermal Master Mix (OptiGene), 0.8 µM each of FIP and BIP, 0.2 µM each F3 and B3 and 1 ng of DNA. All *Acipenser* species (except *A. oxyrinchus*, whose DNA was not available at the time) and *Huso huso* were tested. A negative control was included in the LAMP assay. Reactions were performed on Franklin real time thermocycler (Biomeme) using the isothermal program (3x sample mode) at 65 °C during 1 hour followed by a cooling curve from 95°C to 75 °C (fluorescence reading every 0.5 °C). LAMP products were evaluated based on threshold time T_t_ values (Kubota and Jenkins 2015) and specificity on melting curves.

## Results

### PCR amplification

The DNA from all species amplified the fish-specific marker at the expected size (403 bp or 348 bp) with a weak intensity for *A. sturio* (Figure 1). Only DNA from *A. sturio* amplified the species-specific marker (142 bp). It should be noted that the two DNA quantities tested (2 ng and 20 ng) gave the same results.

**Figure 1:**
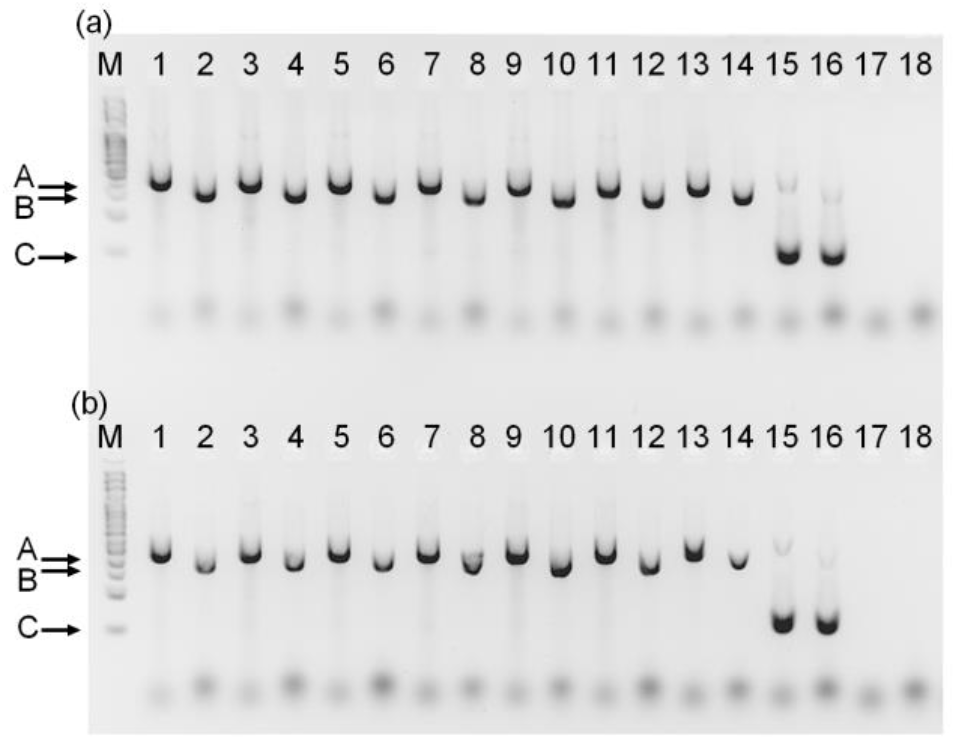
Electrophoretic separation of PCR products from different *Acipenser* species and *Huso huso* with fish specific primers (A bands: L14735/ H15149R pair and B band: L14841/ H15149) pair) and *A. sturio* specific primers (C band) in 3 % agarose gel. M: 100 bp DNA ladder (Solis Biodyne, Estonia), 20 ng (a) or 2 ng (b) of input DNA from: 1 and 2: *A. gueldenstaedtii*, 3 and 4: *A. stellatus*, 5 and 6: *A. transmontanus*, 7 and 8: *A. baerii*, 9 and 10: *A. persicus*, 11 and 12: *Huso huso*, 13 and 14: *A. oxyrinchus*, 15 and 16: *A. sturio*, 17 and 18: water as negative control.

### qPCR amplification

qPCR assays unambiguously detected the specific target species *A. sturio* with C_t_ values of 27.45 ± 0.02 for each 1 ng replicate on the LightCycler. Other species and negative control showed for all triplicates C_t_ values higher than the cut-off value (*i*.*e* total number of qPCR cycles; in this study 40). On the Franklin cycler, 1 ng of *A. sturio* DNA was detected with a C_t_ value of 25 and C_t_ values higher than 40 (i.e. no detectable amplification) for the other species and the negative control. These results demonstrate that the qPCR assay developed in this study is highly *A. sturio* species-specific and does not exhibit amplifications with DNA from other closely related species. Positive detection of DNA quantity down to 1 pg was achieved with a C_t_ value of 35 using both the LightCycler 480 and the field-ready Franklin™ Real-Time PCR Thermocycler.

### LAMP amplification

LAMP melting curve analysis showed a single peak at 89 °C only for *A. sturio* species demonstrating the specificity of the reaction. T_t_ value was 92.2 for *A. sturio* showing efficient amplification, while other species and the negative control showed no amplification (T_t_ value undetermined). Positive detection of DNA quantity down to 100 pg was achieved.

## Discussion

Within the frame of the *A. sturio* restoration programs, incidental captures in the wild happen which allow to know more about the ecology of the species. For example, at sea, those observations allow to assess the species habitat preferences and priority area for conservation can be defined thanks to modelling approaches (e.g. Charbonnel et al. 2023). At sea, at the moment, the risk of confusion between species is low but it is high within the watersheds (freshwater and estuary) where most exotic species are encountered and difficult to morphologically distinguished by non-specialists (Acolas *et al*. 2018). To secure the identification and to be able to use those observations for habitat assessment and improvement of our knowledge of the ecology of the species, quick and easy methods are needed to verify the species. This work is a first step helping the identification for managers and scientists. We can imagine either using fin clips on captured individuals or environmental DNA and associated genetic tools that can be implemented in the field to extend the monitoring to key areas such as spawning areas to confirm the return of the native species with limited species confusion risk.

To our knowledge, this study is the first to report PCR-based assays and LAMP technique for the detection of the European sturgeon *A. sturio*. The qPCR reaction took only one hour, which was comparable to real-time LAMP. PCR had a reaction time of 1 h 25 min with an additional time of nearly 1h for agarose gel electrophoresis, making LAMP and qPCR the quickest methods compared with conventional PCR. However, the LAMP assay implemented in this study was four times more expensive than the qPCR making the latter a method of choice for fast and affordable *A. sturio* identification. Moreover, qPCR detection showed the higher sensitivity compared to LAMP, which make the method more appropriate for environmental DNA approaches where low DNA quantity is expected. Both qPCR and LAMP can be performed in the field using Biomeme’s Franklin™ Real-Time PCR Thermocycler. Nevertheless, the identification of *A. sturio* by PCR remains the most reliable method because of the use of a fish-specific marker as a positive control validating negative results from the species-specific marker. The use of a positive control in qPCR would require the design of a fish-specific probe labeled with a different reporter dye that the one used to detect *A. sturio* at an additional cost and experiment complexity. PCR followed by agarose gel is also the most accessible to any laboratory or field application (using for instance portable Bento Lab) and the analysis of the result is visual and easy to understand and reproducible. Depending on the application, the three developed methods have their benefits in term of cost, sensitivity and ease of use, but all can be implemented in the field to specifically detect the native *A. sturio* from other non-native closely-related species.

## Acknowledgments

Technical developments and laboratory experiments were performed at the PGTB (doi:10.15454/1.5572396583599417E12) with the help of Christophe Boury, Marie Massot and Erwan Guichoux, and contributions from Héloïse Delval and Léni Guillotin. We wish to thank Antoine Mouilleron from the hatchery “Huso” (Neuvic, South West of France) who kindly provided the fin samples for *A. gueldenstaedtii, A. stellatus, A. transmontanus, A. baerii, A. persicus*, and *Huso huso* with the help of P. Chèvre and R. le Barh (INRAE). The *A. sturio* samples comes from the French captive stock (Saint Seurin sur l’Isle, France). We wish to thank Joern Gessner (IGB, Berlin) who provided fin tissue of *A. oxyrinchus* from the German captive stock. This study has been carried out with financial support from the French National Research Agency (ANR) in the frame of the Investments for the Future Programme, within the Cluster of Excellence COTE (ANR-10-LABX-45).

## Notes

Funding statement: This study has been carried out with financial support from the French National Research Agency (ANR) in the frame of the Investments for the Future Programme, within the Cluster of Excellence COTE (ANR-10-LABX-45).

Conflict of interest: The authors declare that they have no conflict of interests.

### Competing Interest Statement

The authors have declared no competing interest.

