## supporting material for "Species-level identification of the critically endangered European sturgeon *Acipencer sturio* using DNA-based approaches"

**Table S1:** Accession number of cytochrome b gene used for PCR and qPCR primer design.

| Species | GenBank accession number |
| --- | --- |
| *Huso huso* | AY846703.1; AY846702.1; FJ974044.1; KC130117.1; KC130116.1; KC130115.1; KC130114.1; KC130113.1; KC130112.1; KC130111.1; KC130110.1; AJ245840.1; AY846712.1; AY846711.1; AY846710.1; AY846709.1; AY846708.1; AY846707.1; AY846706.1; AY846705.1; AY846704.1; |
| *Acipenser baerii* | EU733253.1; AJ245825.1; FJ010862.1; FJ010861.1; FJ010860.1; |
| *Acipenser persicus* | AJ245837.1; AJ245836.1; AJ245835.1; |
| *Acipenser gueldenstaedtii* | EU733252.1; JN698961.1; AJ277594.1; AJ277597.1; AJ277596.1; AJ277595.1; FJ974040.1; KC130102.1; KC130101.1; KC130100.1; KC130099.1; KC130098.1; KC130097.1; KC130096.1; KC130095.1; KC130094.1; KC130093.1; KC130092.1; KC130091.1; KC130090.1; AJ563396.1; AJ563395.1; AJ563394.1; AJ563393.1; AJ563392.1; AJ563391.1; AJ563390.1; AJ563389.1; AJ563388.1; AJ563387.1; AJ563386.1; AJ563385.1; AJ249692.1; AJ245827.1; AJ245826.1; |
| *Acipenser stellatus* | FJ974042.1; KC130109.1; KC130108.1; KC130107.1; KC130106.1; KC130105.1; KC130104.1; KC130103.1; AJ249693.1; AY846701.1; AY846700.1; AY846699.1; AY846698.1; AY846697.1; AY846696.1; AY846695.1; AY846694.1; AY846693.1; AY846692.1; AY846691.1; AY846690.1; AY846689.1; AY846688.1; AY846687.1; AY846686.1; AY846685.1; AY846684.1; AY846683.1; AY846682.1; AY846681.1; AY846680.1; |
| *Acipenser sturio* | EU733250.1; EU733249.1; EU733248.1; FJ974043.1; AJ428497.1; |
| *Acipenser transmontanus* | KF013247.1 |

**Table S2**: LAMP primer sequences and characteristics (from PrimerExplorer v5). TTTTTT represents the linker between primers of F1c and F2 (FIP) and those of B1c and B2 (BIP).

| Label | 5' position | 3' position | Length | Tm | 5'dG | 3'dG | GC  rate | Sequence |
| --- | --- | --- | --- | --- | --- | --- | --- | --- |
| F3 | 284 | 301 | 18 | 59.7 | -3.77 | -6.10 | 0.61 | CTTTTGGCCTCCTCTGGC |
| B3 | 472 | 492 | 21 | 59.2 | -4.76 | -6.57 | 0.52 | GTCTGATACTGGGATACTGCG |
| FIP (F1c+F2) |  |  | 38 |  |  |  |  | AGGCTCCTGCATGGGCTAGGTTTTTTCTGGCACAGGATGAACCG |
| BIP (B1c+B2) |  |  | 43 |  |  |  |  | CCTTCACCTGGCTGGGGTATCTTTTTTGGGGGCTTCATGTTAATGATTG |

**Figure S1**: DNA sequences alignment of the *A. sturio* specific primers (first row) with the corresponding portion of cytochrome b gene in *Acipenser* and *Huso* species.

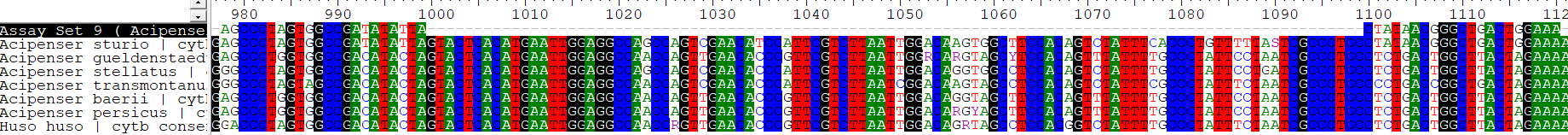

Table S2: Accession number of COI gene used for LAMP primer design.

| Species | GenBank accession number |
| --- | --- |
| *Acipenser sturio* | NC_027417.1; KP997216.1; KJ552473.1; KJ552406.1; |
| *Acipenser stellatus* | HQ960589.1; HQ960588.1; HQ960587.1; HQ960586.1; HQ960585.1; KC500133.1; KC500132.1; KC500131.1; KC500130.1; KC500129.1; KC500128.1; KC500127.1; KC500126.1; KC500125.1; KC500124.1; KC500123.1; KC500122.1; KC500121.1; KC500120.1; KC500119.1; MK903722.1; FJ809739.1; FJ809738.1; FJ809737.1; FJ809736.1; FJ809735.1; JQ623906.1; KC578824.1; KC578823.1; AJ585050.1; |
| *Acipenser gueldenstaedtii* | HQ960593.1; HQ960592.1; HQ960591.1; HQ960590.1; KM286425.1; KM286424.1; KM286423.1; KC500103.1; KC500102.1; KC500101.1; KC500100.1; KC500099.1; KC500098.1; KC500097.1; KC500096.1; KC500095.1; KC500094.1; KC500093.1; KC500092.1; KC500091.1; KC500090.1; KC500089.1; KC500088.1; MK903725.1; MK903723.1; FJ809729.1; FJ809728.1; FJ809727.1; FJ809726.1; FJ809725.1; JQ623904.1; KC578837.1; KC578836.1; KC578835.1; |
| *Acipenser persicus* | FJ809724.1; FJ809723.1; FJ809722.1; FJ809721.1; FJ809720.1; |
| *Acipenser baerii* | KM286422.1; KM286421.1; KM286420.1; GQ328816.1; GQ328815.1; GQ328814.1; GQ328813.1; GQ328812.1; GQ328811.1; GQ328810.1; GQ328809.1; GQ328808.1; GQ328807.1; GQ328806.1; GQ328805.1; GQ328804.1; GQ328803.1; GQ328802.1; GQ328801.1; GQ328800.1; GQ328799.1; GQ328798.1; GQ328797.1; GQ328796.1; GQ328795.1; GQ328794.1; GQ328793.1; GQ328792.1; GQ328791.1; GQ328790.1; GQ328789.1; GQ328788.1; GQ328787.1; GQ328786.1; GQ328785.1; GQ328784.1; GQ328783.1; MK903726.1; MK903724.1; KC578844.1; GU804895.1; FJ205563.1; FJ205562.1; FJ205561.1; FJ205560.1; KP833627.1; KP833626.1; KP833625.1; KP833624.1; KP833623.1; KP833622.1; KP833621.1; KP833620.1; KP833619.1; KP833618.1; KP833617.1; KC578845.1; KC578843.1; KC578842.1; KC578836.1; HM902201.1; HM902200.1; |
| *Huso huso* | MK903721.1; FJ809719.1; FJ809718.1; FJ809717.1; FJ809716.1; FJ809715.1; |
| *Acipenser transmontanus* | KX145032.1; MK903720.1; MK903719.1; MK903717.1; KF558304.1; EU523891.1; EU523890.1; EU523889.1; EU523888.1; EU523887.1; AB042837.1; |

**Figure S2**: LAMP primer designed for *Acipenser sturio* COI gene. (A) The location of *A. sturio* COI-Design50 in the partial COI gene sequence (from PrimerExplorer v5). (B) Schematic representation of the LAMP primers used in this study (Note that FIP and BIP primers are made of F1c+F2 and B1c+B2 respectively).

(A)

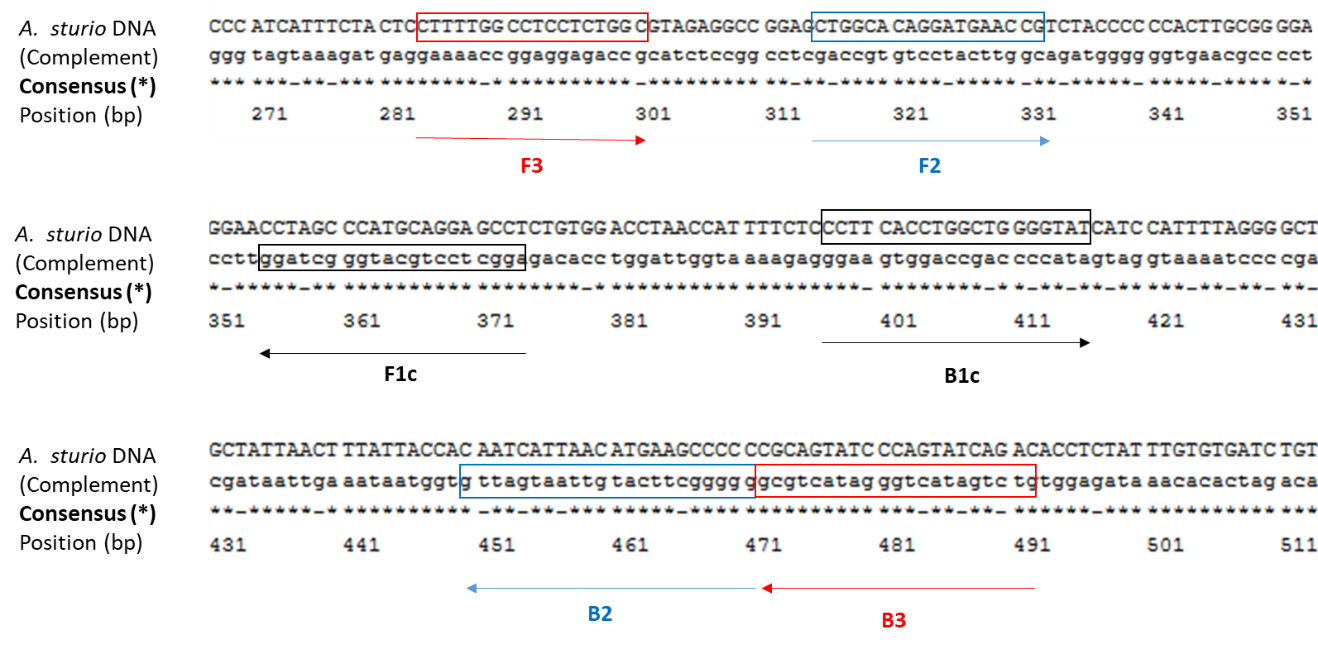

(B)

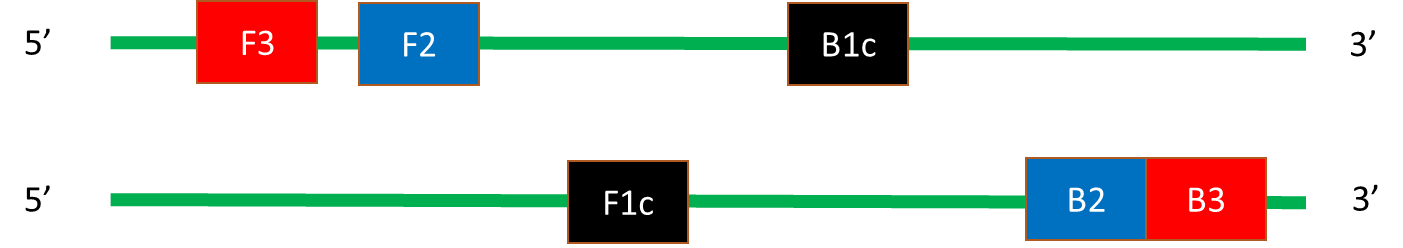
